# Mapping the Role of AcrAB-TolC Efflux Pumps in the Evolution of Antibiotic Resistance Reveals Near-MIC Treatments Facilitate Resistance Acquisition

**DOI:** 10.1101/2020.07.07.192856

**Authors:** Ariel M. Langevin, Imane El Meouche, Mary J. Dunlop

## Abstract

Antibiotic resistance has become a major public health concern as bacteria evolve to evade drugs, leading to recurring infections and a decrease in antibiotic efficacy. Systematic efforts have revealed mechanisms involved in resistance; yet, in many cases, how these specific mechanisms accelerate or slow the evolution of resistance remains unclear. Here, we conducted a systematic study of the impact of the AcrAB-TolC efflux pump on the evolution of antibiotic resistance. We mapped how population growth rate and resistance change over time as a function of both the antibiotic concentration and the parent strain’s genetic background. We compared the wild type strain to a strain overexpressing AcrAB-TolC pumps and a strain lacking functional pumps. In all cases, resistance emerged when cultures were treated with chloramphenicol concentrations near the MIC of their respective parent strain. The genetic background of the parent strain also influenced resistance acquisition. The wild type strain evolved resistance within 24 h through mutations in the *acrAB* operon and its associated regulators. Meanwhile, the strain overexpressing AcrAB-TolC evolved resistance more slowly than the wild type strain; this strain achieved resistance in part through point mutations in *acrB* and the *acrAB* promoter. Surprisingly, the strain without functional AcrAB-TolC efflux pumps still gained resistance, which it achieved through upregulation of redundant efflux pumps. Overall, our results suggest that treatment conditions just above the MIC pose the largest risk for the evolution of resistance and that AcrAB-TolC efflux pumps impact the pathway by which chloramphenicol resistance is achieved.

**IMPORTANCE:** Combatting the rise of antibiotic resistance is a significant challenge. Efflux pumps are an important contributor to drug resistance; they exist across many cell types and can export numerous classes of antibiotics. Cells can regulate pump expression to maintain low intracellular drug concentrations. Here, we explored how resistance emerged depending on the antibiotic concentration, as well as the presence of efflux pumps and their regulators. We found that treatments near antibiotic concentrations that inhibit the parent strain’s growth were most likely to promote resistance. While wild type, pump overexpression, and pump knock out strains were all able to evolve resistance, they differed in the absolute level of resistance evolved, the speed at which they achieved resistance, and the genetic pathways involved. These results indicate that specific treatment regimens may be especially problematic for the evolution of resistance and that the strain background can influence how resistance is achieved.

## INTRODUCTION

Despite the new wave of antibiotic discovery (1–5), bacteria continue to acquire resistance shortly after the introduction of new drugs for medicinal and industrial applications (6, 7). This is due in large part to the overuse of antibiotics, which results in pressures that drive resistance (8). With limited novel antibiotics and numerous futile antibiotics, doctors and scientists alike are presented with the challenge of how to best treat infections while keeping the evolution of resistance in check.

Adaptive evolution studies have begun exploring how certain antibiotic pressures influence the evolution of resistance. For instance, studies using a ‘morbidostat’—a continuous culture device that dynamically adjusts antibiotic concentrations to inhibitory levels—have found numerous targets that can be readily mutated to promote resistance (9–11), as well as identifying how drug switching can limit the evolution of resistance (12). While these studies have provided pivotal insights for this field, the morbidostat design causes antibiotic concentrations to rise to levels that exceed clinically relevant concentrations due to toxicity for patients (13). In recognition of the drug concentration-dependent nature of evolution, researchers have begun to explore bacterial evolution under treatment conditions with lower antibiotic concentrations as well. Wistrand-Yuen *et al*. found that bacteria grown in subinhibitory drug concentractions were still able to achieve high levels of resistance (14–16). Notably, the study identified that the same antibiotic produced unique evolutionary pathways when cells were treated with subinhibitory concentrations as opposed to inhibitory concentrations (14).

One limitation of current studies within the field is that they can be difficult to compare due to variations in experimental parameters, such as species, antibiotics, or other experimental conditions (17). Given the unique evolutionary pathways at different antibiotic concentrations, systematic mapping of these evolutionary landscapes could provide an improved understanding of which conditions pose the highest risk by allowing direct comparisons between different antibiotic concentrations. For instance, Jahn *et al*. demonstrated that variations in treatment dynamics can significantly alter evolved resistance for some antibiotics, such as tetracycline, but not others, such as amikacin and piperacillin (18). Other evolution experiments that were systematically conducted using a range of concentrations for beta-lactams (19) and erythromycin (20) have highlighted the concentration-dependent adaptability of *Escherichia coli*.

There are many mechanisms by which antibiotic resistance can be achieved, including enzymatic inactivation, alteration of antibiotic binding sites, and increased efflux or reduced influx of antibiotics (21, 22). Efflux pumps are omnipresent in prokaryotic and eukaryotic cells alike, and are an important contributor to multi-drug resistance (23). AcrAB-TolC in *E. coli* is a canonical example of a multi-drug efflux pump, providing broad-spectrum resistance and raising the MIC of at least nine different classes of antibiotics (24). The pump is composed of three types of proteins: the outer membrane channel protein, TolC; the periplasmic linker protein, AcrA; and the inner membrane protein responsible for substrate recognition and export, AcrB (23). Using the proton motive force, AcrB actively exports antibiotics from the cell (23, 25). The presence of AcrAB-TolC efflux pumps can increase a strain’s MIC from ~2-fold to ~10-fold, depending on the antibiotic (26–28). Furthermore, genes associated with these multi-drug resistant efflux pumps, including their local and global regulators, are common targets for mutation as strains evolve high levels of drug resistance (15, 29–32).

Recent studies have indicated that in addition to providing modest increases in the MIC due to drug export, pumps can also impact mutation rate and evolvability of strains, which may ultimately be more important for the acquisition of high levels of drug resistance. For example, Singh *et al*. found that mutants overexpressing *acrAB* emerged first and afterwards these mutants could evolve high levels of quinolone resistance (33). In addition, heterogeneity in efflux pump expression can predispose subsets of bacterial populations with elevated *acrAB* expression to mutation even prior to antibiotic treatment (34). Deleting genes associated with efflux pumps, such as *tolC*, can also reduce evolvability under antibiotic exposure (35). Further, a recent study in *Staphylococcos aureus* found that higher NorA pump levels increased evolvability, and that adding a pump inhibitor could prevent resistance evolution (36). These studies provoke the question of how AcrAB-TolC efflux pumps impact the evolution of drug resistance.

Our overall goal in this study was to identify how strains with different AcrAB-TolC genotypes evolve antibiotic resistance over time under a range of chloramphenicol concentrations. Chloramphenicol is both a well-validated substrate of AcrAB-TolC and can serve as a last resort antibiotic in multi-drug resistant infections, as most clinical isolates are still susceptible to this drug (37, 38). To identify how AcrAB-TolC impacts the evolution of resistance, we used a turbidostat as an evolutionary platform (39) and measured changes in fitness and resistance. We evolved three strains with different levels of AcrAB-TolC: a wild type strain with the native regulatory network controlling AcrAB-TolC expression (WT); a strain which lacks the local regulator AcrR (AcrAB+), which results in a 1.5 to 6-fold increase in expression of the pumps (40–42); and a strain lacking functional AcrAB-TolC efflux pumps (Δ*acrB*). We allowed the cultures to grow and evolve for 72 h in continuous culture while continuously recording growth rates. We periodically sampled the cultures and assessed the population’s resistance. We then charted the evolutionary landscapes for each of the three strains under different chloramphenicol concentrations to identify which circumstances gave rise to resistance.

## RESULTS

In order to systematically evaluate the evolutionary landscape of efflux pump-mediated antibiotic resistance, we used the eVOLVER, a modular turbidostat capable of growing independent cultures in parallel (39). This platform allowed us to track a culture’s fitness by measuring growth rate continuously over multi-day experiments. In addition to this, we collected samples at selected intervals and, with these samples, performed antibiotic disc diffusion assays to assess the population’s resistance and spot assays to quantify the presences of high-resistance isolates within the population (Figure 1).

**Figure 1.**
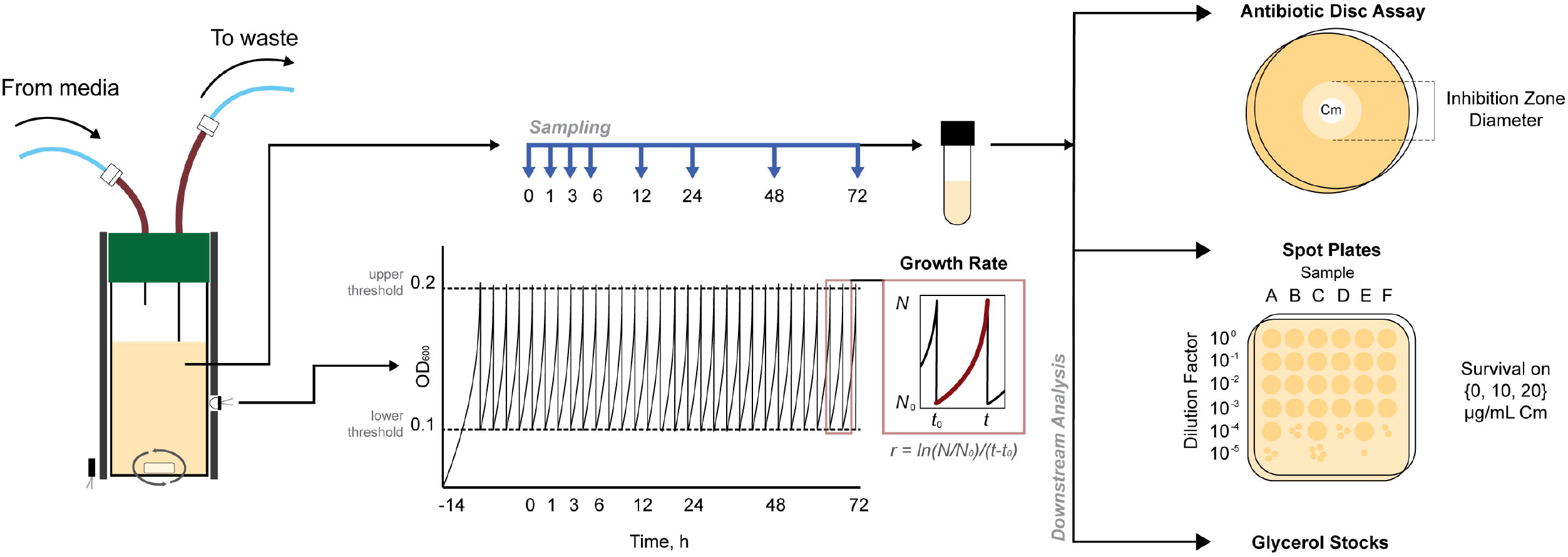
Evolution experiment schematic. We used the eVOLVER, a modular turbidostat, as an evolutionary platform to measure and record absorbance data at 600 nm (OD_600_). We calculated growth rate after each dilution event and collected samples at defined timepoints (t = 0, 1, 3, 6, 12, 24, 48, 72 h). We performed antibiotic disc assays and spot plate assays for all samples.

We mapped growth rates over time for cultures subjected to a range of chloramphenicol treatment concentrations (Figure 2A & Figure S1). To compare across strains, we defined MIC^0^_parent_ as the MIC of the parent strain (MIC^0^_WT_ = 2 μg/mL, MIC^0^_AcrAB+_ = 2 μg/mL, MIC^0^_Δ*acrB*_ = 0.5 μg/mL). We found similar values for MIC^0^_WT_ and MIC^0^_AcrAB+_ (Figure S2), which may be due to induction of efflux pump expression in the WT strain in the presence of chloramphenicol. Prior studies have shown that the presence of stress can increase pump expression by 4-fold (40, 43), which is comparable to the impact of deleting *acrR* (40–42). We found that treatment with high concentrations of chloramphenicol repressed bacterial growth for multiple days. We observed this growth inhibition at ~10 μg/mL for WT and AcrAB+, and at ~2 μg/mL for Δ*acrB*. These inhibitory concentrations represent treatments of ~5x MIC^0^_parent_ for all three strains. We found that cultures grown in lower chloramphenicol concentrations were able to recover growth. For example, when we treated cultures with ~1-2x MIC^0^_parent_, we observed a significant decrease in the growth rate between 0 and 12 h (Table S1). However, after 12 to 24 h, growth in these populations was partially restored. At lower treatment concentrations (<1x MIC^0^_parent_), all cultures were able to grow, though usually at a deficit compared to the 0 μg/mL chloramphenicol condition. For all three strains, there were qualitatively similar growth recovery patterns, with an initial growth repression phase followed by a partially restored growth phase (Figure S1).

**Figure 2.**
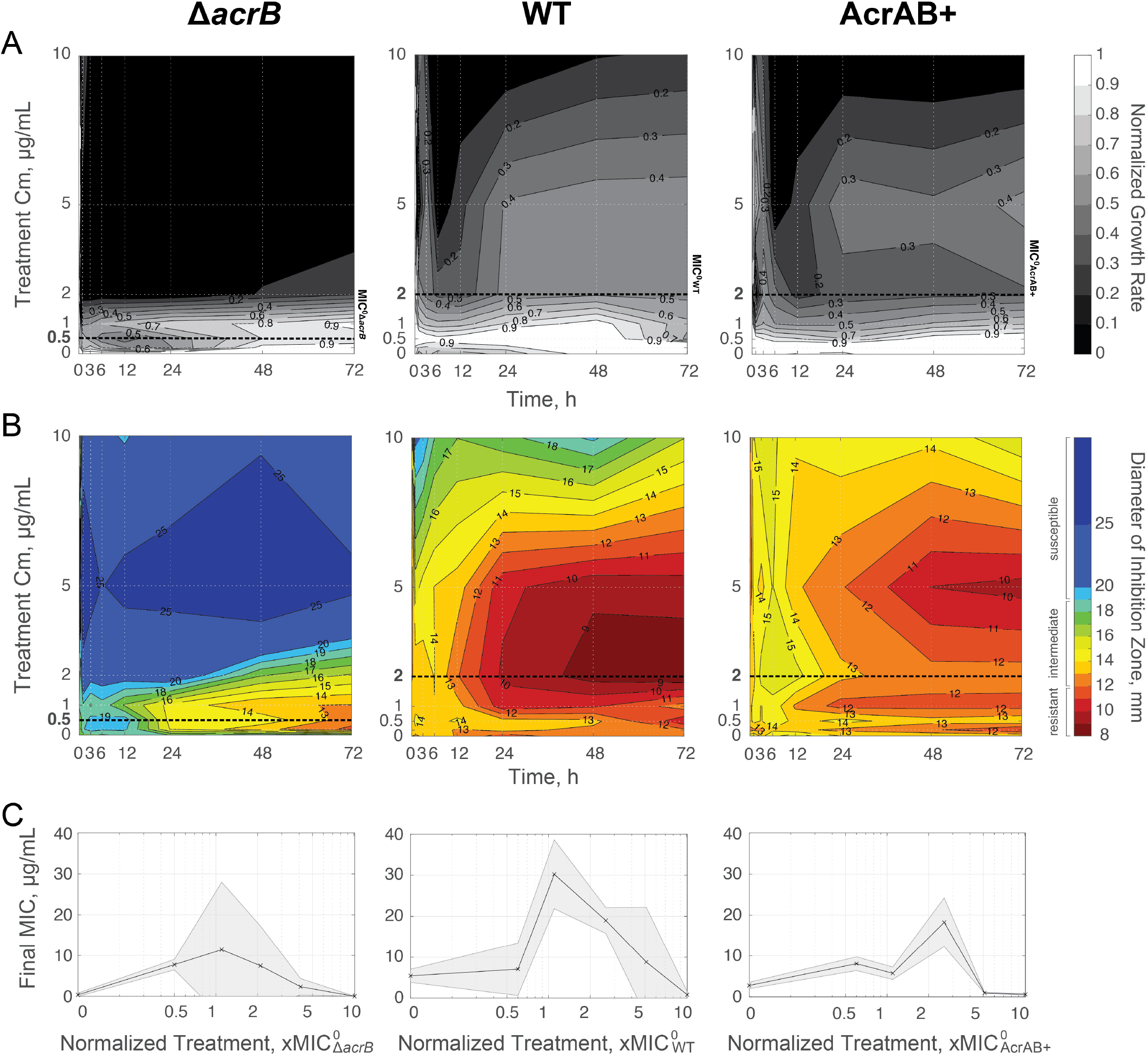
Temporal landscapes based on treatment concentration of chloramphenicol. **(A)** Average growth rate. Growth rates are normalized to growth of strains at t = 0 h; for raw data see Figure S1. Lighter areas represent growth rates closer to pre-treatment values; darker areas represent reduced growth rates. MIC^0^_parent_ concentration is denoted with a bold dashed line for each strain (Figure S2). **(B)** Average resistance. Diameter of inhibition zones were plotted for each time and treatment. Smaller inhibition zones are shown in red and correspond to resistant cells (£12 mm) and larger inhibition zones are shown in blue and represent susceptible cells (≥19 mm); intermediate inhibition is shown with color scale from orange to green. MIC^0^_parent_ is denoted with a bold dashed line. **(C)** Final resistance at 72 h based on treatment concentration normalized to MIC^0^_parent_. The final, absolute MIC is calculated based on data from Figure S5. Data points show the mean of three biological replicates. Shaded error bars show standard deviation.

The growth rate results suggested the evolution of drug resistance within the population (9, 18). To quantify this, we used an antibiotic disc assay to map the corresponding resistance levels (Figure 2B & Figure S3). We found distinct increases in resistance levels that corresponded to populations which recovered growth. While there were qualitative similarities for the three strains, the timing and level of resistance achieved was dependent on the strain background. We classified populations as resistant when their inhibition zone diameters were smaller than 12 mm, following established standards for antimicrobial susceptibility testing (44). The WT strain gained resistance under a broad range of chloramphenicol treatment concentrations; this resistance emerged within 24 h when cells were treated with ~1-2x MIC^0^_WT_. The AcrAB+ strain, where efflux pumps are overexpressed, was able to evolve resistance as well, albeit at a slower rate and at lower levels than WT. AcrAB+ achieved resistance within 48 h when treated with 2.5x MIC^0^_AcrAB+_, but the range of chloramphenicol concentrations that resulted in resistance was narrower than for the WT strain. The Δ*acrB* cells achieved resistance more slowly, but for the range of ~1-2x MIC^0^_Δ*acrB*_ chloramphenicol cultures were still able to reach resistant levels (Figure 2B & Figure S3).

To compare the ultimate evolved resistance levels, we calculated the final, absolute MIC of the populations at 72 h. When we normalized the treatment concentration by MIC^0^_parent_, we found that treatments concentrations ~1-2x MIC^0^_parent_ evolved the most resistant populations (Figure 2C). Selective pressures of subinhibitory antibiotic concentrations have often been considered high-risk for the evolution of resistance (14, 45). Yet, our results indicated that concentrations near or just above MIC^0^_parent_ lead to the highest resistance levels in these conditions. In short, all three strains were able to evolve resistance when treated with ~1-2x MIC^^0^^parent chloramphenicol, with WT achieving the highest final, absolute MIC of the three strains. WT evolved more rapidly than AcrAB+ or Δ*acrB*. Moreover, the relative range of chloramphenicol concentrations that supported the evolution of resistance in the AcrAB+ strain was narrower than for WT or Δ*acrB* strains.

We next asked how resistance and growth changed through time. We found that in the absence of antibiotics, the trajectories trended largely towards faster growth, with minimal changes to resistance levels (Figure 3). With subinhibitory chloramphenicol treatments, we observed that the populations first experienced a slight growth decrease, followed by increased resistance, and then restored growth within 48 h. While these populations did gain resistance, they did not tend to reach very high final MIC values in absolute terms, with inhibition zone diameters just at the border of being defined as resistant. In contrast, with inhibitory chloramphenicol treatment, there was a more dramatic reduction in growth within the first 12 h. Though growth was impacted, the populations tended to walk towards high resistance during this period. As depicted in the schematics, the zig-zag patterns trending towards high resistance may be indicative of the cultures acquiring resistant mutations and compensating for the associated fitness costs of these mutations. Finally, at high chloramphenicol concentrations, bacteria first became more susceptible and then stopped growing entirely within 12 h; growth was never restored for these populations. We found that all strains followed similar evolutionary trajectories while balancing the trade-off between growth and resistance. These findings highlight the importance of using antibiotic concentrations that are sufficiently inhibitory.

**Figure 3.**
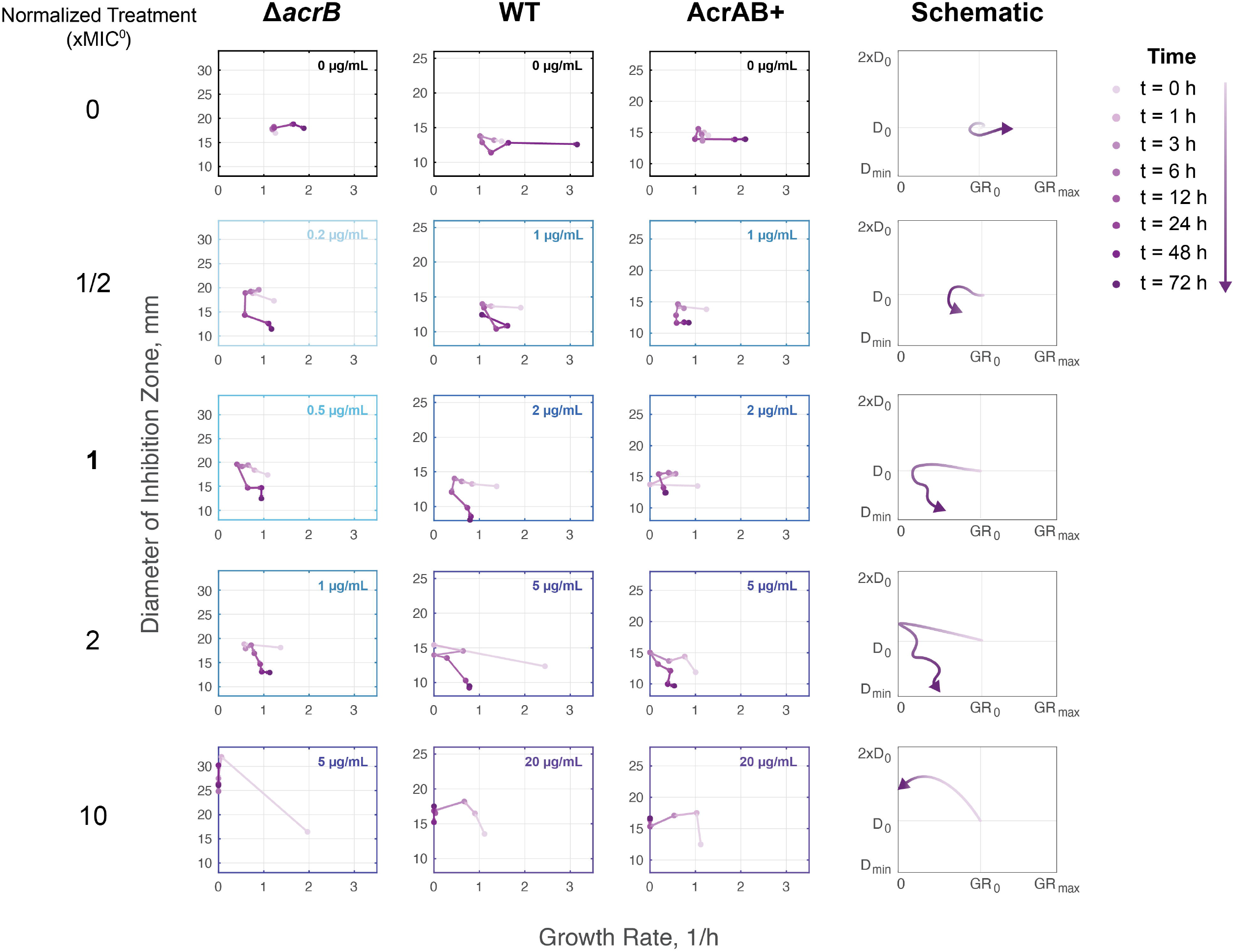
Resistance and Fitness Evolution Trajectories. **(A)** Average diameter of inhibition zone and average growth rate plotted against each other. Lighter purple markers represent trajectories occurring earlier; darker purple are later timepoints. The longer the distance between markers, the greater the change between time points. Colors of boxes indicate the absolute treatment concentration for the depicted trajectories. **(B)** Schematics summarize patterns for each treatment concentration (xMIC^0^_parent_). Schematic plots show growth rate in terms of initial growth rate (GR_0_) and maximum physiological growth rate (GR_max_). Resistance is shown in terms of relative diameter of inhibition, where D_0_ is the diameter of inhibition at t = 0 h and D_min_ is the diameter of the antibiotic disc.

While these results tell us about the growth rate and resistance of the overall population, it is difficult to determine if sub-populations of cells within the culture have acquired high levels of resistance from disc assays alone. First, because the disc assays do not quantify resistance associated with individual cells in the culture, they cannot reveal the presence of sub-populations of resistant and susceptible cells. Second, beyond a certain resistance level, cells will grow up to the boundary of the disc; thus, it is not possible to quantify resistance increases beyond this. Determining which conditions can give rise to high levels of resistance is important for revealing particularly dangerous treatment regimes. In addition, sub-populations with increased resistance to one antibiotic can promote cross-resistance to other drugs (45).

To quantify the fraction of resistant cells that emerged during our evolution experiment, we conducted a spot assay, in which we measured the fraction of the population capable of surviving on specific chloramphenicol concentrations. For all three strains, we observed sub-populations that were capable of growing on 10 μg/mL chloramphenicol (Figure 4A & Figure S4). Interestingly, these cells primarily emerged from treatment conditions with lower levels of chloramphenicol, and not from conditions where cells were subjected to 10 μg/mL chloramphenicol. For example, at least 0.1% of the population from each of the three WT replicates that were treated at 2 μg/mL chloramphenicol could survive on 10 μg/mL at the end of the experiment. We did find cases where WT cells treated with 10 μg/mL evolved resistance to 10 μg/mL, however this was less common than in lower treatment concentrations. Thus, cultures were able to evolve resistance to higher levels of chloramphenicol than they were subjected to, a feature that was most pronounced when treatments were just above or at MIC^0^_WT_. These results closely match trends in the population’s overall resistance (Figure 2B). We also found isolates capable of growing on 20 μg/mL chloramphenicol, with a reduced frequency relative to 10 μg/mL (Figure 4B & Figure S4).

**Figure 4.**
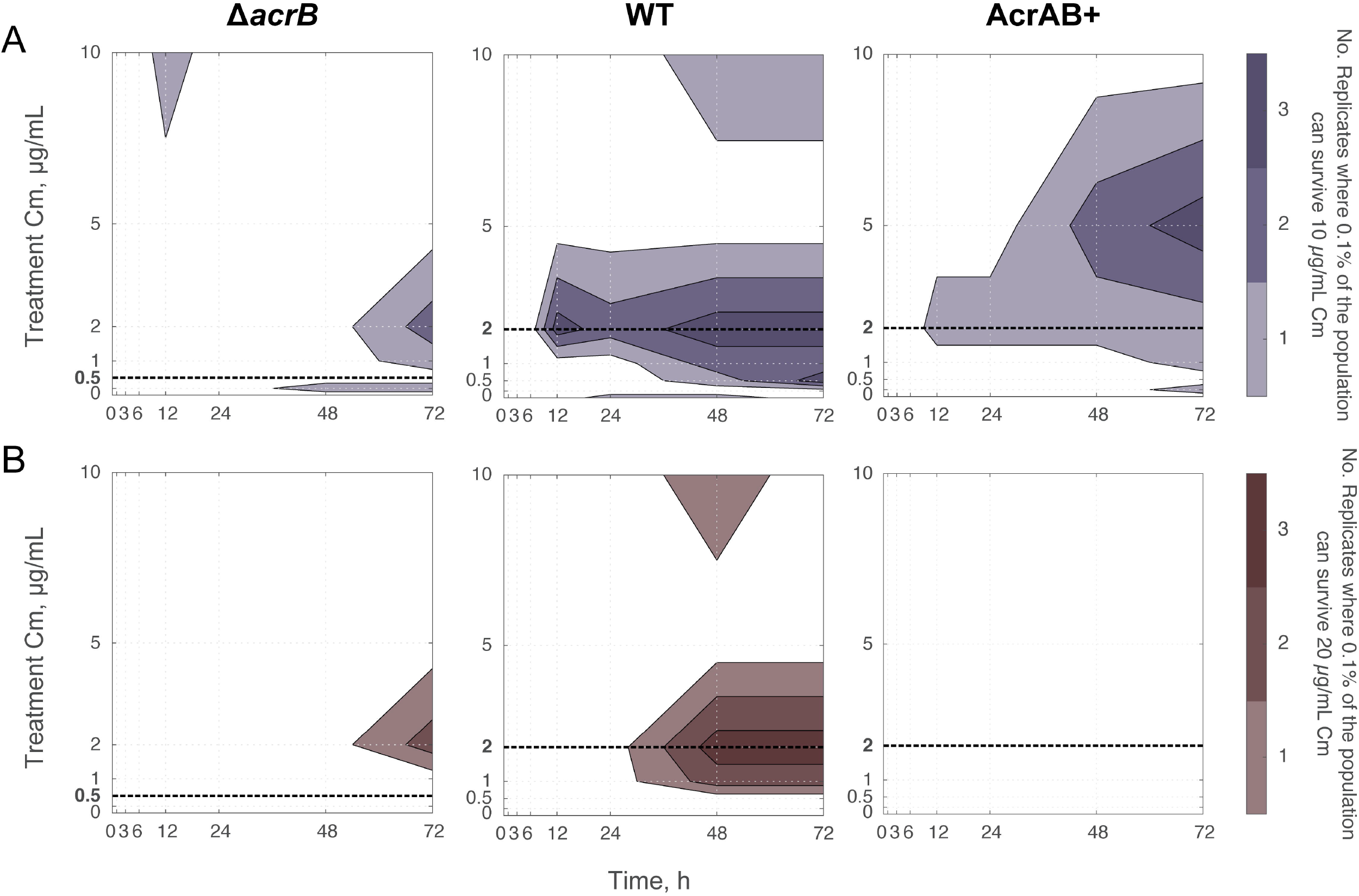
Number of Biological Replicates with Highly Resistant Sub-populations through Time. Number of biological replicates that had a subpopulation greater than 0.1% of their total population, which could grow on LB plates containing **(A)** 10 μg/mL or **(B)** 20 μg/mL chloramphenicol. Raw data are shown in Figure S4. Initial populations contained ~10^7^ CFUs. MIC^0^_parent_ is denoted with a bold dashed line.

In contrast, the AcrAB+ strain was capable of evolving resistance to 10 μg/mL when treated with 5 μg/mL chloramphenicol; yet, surprisingly, AcrAB+ never produced a sub-population that was able to grow on 20 μg/mL as the WT did. Meanwhile, despite the higher initial susceptibility of Δ*acrB* (MIC^0^_Δ*acrB*_ < MIC^0^_WT_ and MIC^0^_AcrAB+_), the Δ*acrB* strain consistently produced subpopulations that were able to grow at 20 μg/mL chloramphenicol by 72 h. This sub-population appeared for chloramphenicol concentrations around 2 μg/mL, similar to the WT strain.

A key question remained: which mutations were responsible for the increases in resistance we observed? To address this, we used whole genome sequencing to analyze three biological replicates from the 72 h timepoint for the WT, AcrAB+, and Δ*acrB* strains (Table S2). For the WT strain, each of the sequenced isolates contained a single point mutation in the DNA binding region of *marR*, which can upregulate AcrAB-TolC efflux pumps and expression of other stress response genes (46). Two of these point mutations were missense mutations in *marR* and have been observed in other studies (47–51). Additionally, one isolate had a missense mutation in the periplasmic encoding region of *acrB*. The other two isolates had an IS1 or IS5 insertional sequence interrupting *acrR*, which is known to upregulate *acrAB* (52). One question these results raise is why the AcrAB+ strain, where *acrR* is removed, is outperformed by WT strains with mutations in *acrR*. A potential explanation for this is that the ‘marbox’ through which *acrAB* is upregulated sits within *acrR* (53). The AcrAB+ strain lacks this marbox (54), while in the sequenced isolates the insertion sequence is located further upstream in *acrR* and the marbox remains intact, providing global stress response regulation while eliminating the impact of the local repressor. Thus, the exact position of the insertion sequence matters. These sequencing results indicate that strains containing AcrAB-TolC efflux pumps use mutations related to the pumps and their regulation to optimize survival and increase resistance in the presence of chloramphenicol.

When we evolved the AcrAB+ strain and performed whole genome sequencing of the most resistant isolates, all isolates had mutations in the noncoding, promoter region of *acrAB* (Table S2). These mutations indicate that the AcrAB+ strain might require further tuning of *acrAB* expression for improved resistance. Further, two of these isolates also had missense mutations in the coding region of *acrB* as well. Of these, the V139F missense mutation is known to produce high levels of multidrug resistance by accelerating export for a number of AcrAB-TolC substrates (11, 18, 55, 56). We observed *acrB* Q569L evolve from two different parent strains, WT and AcrAB+, suggesting it plays a role in chloramphenicol export. Additionally, the evolved AcrAB+ isolates all had other mutations less directly related to the AcrAB-TolC efflux pump and its regulators, such as genes related to transcription (*rpoB, yhjB*), fimbriae assembly (*fimD*), or degradation (*clpX*) (Table S2).

In contrast, when we evolved the Δ*acrB* strain, we found that all three isolates had an insertion sequence located in *acrS* (Table S2). AcrS is the local regulator of the AcrEF-TolC efflux pump, a homolog to AcrAB-TolC (57). This result agrees with findings from Cudkowicz & Schuldiner, who showed that the Δ*acrB* strain gained high resistance by upregulating redundant efflux pumps in *E. coli*, such as AcrEF-TolC or MdtEF-TolC (11). One of the three isolates also contained a missense mutation in the tRNA for selenocysteine (*selA*) and a short insertion sequence in the 16S rRNA of the 30S subunit (*rrsG*), though whether or how these play a role in chloramphenicol resistance is unclear.

## DISCUSSION

In this work, we identified that treating strains with antibiotic concentrations close to MIC^0^parent promotes the evolution of resistance; however, the evolvability and ultimate resistance level achieved differed between WT, AcrAB+, and Δ*acrB* strains. WT populations evolved mutations that conferred high levels of resistance within 24 h after antibiotic exposure. Maximal resistance was evolved at ~1x MIC^0^_WT_, however 0.25-2.5x MIC^0^_WT_ chloramphenicol treatment concentrations all gave rise to resistance. In contrast, AcrAB+ evolved resistance, but this was only possible at precise chloramphenicol concentrations at 2.5x MIC^0^_ActAB+_. The evolved AcrAB+ populations were less resistant than their WT counterparts, and spot assays determining resistance confirmed this trend. In contrast, the Δ*acrB* strain was able to evolve resistance under 1-4x MIC^0^Δ*acrB* chloramphenicol treatments, and ultimately achieved absolute resistance levels comparable to those observed in the WT strain.

Our results identify that antibiotic treatments near MIC^^0^^_parent_ are especially prone to evolving resistance. Reding *et al*. observed this hotspot for adaptability of *E. coli* in the presence of another antibiotic, erythromycin, just below the MIC of their parent strains (20). While doctors measure resistance of bacterial infections, they sometimes prescribe antibiotic treatment prior to obtaining the results of this assay (58) or use a treatment concentration too low to effectively penetrate the infection site (59). This blind treatment could lead to increased levels of resistance (60, 61). These results highlight the presence of regimes that are especially problematic and which should be avoided to limit the evolution of antibiotic resistance.

While we observed that all strains were capable of evolving resistance, sequencing revealed the different pathways that each strain took to achieve this. WT achieved resistance through mutations and insertion sequences in the regulators AcrR and MarR, suggesting that WT cells can fine-tune expression of the AcrAB-TolC pumps to gain resistance to chloramphenicol. Interestingly, these mutations may produce cross-resistance to other antibiotics as well since these regulators control many genes involved in multi-drug resistance (62, 63). AcrAB+ cells utilized mutations in *acrB* and the promoter region controlling its expression to achieve resistance. Δ*acrB* populations achieved resistance by targeting homologous efflux pump systems, such as AcrEF-TolC. Although resistance was slow to emerge in this strain compared to WT or AcrAB+, this alternative pathway for achieving resistance ultimately resulted in levels comparable to those achieved by the WT strain. By charting evolutionary landscapes across different antibiotic concentrations, we have gained insight into treatments that impact the emergence of antibiotic resistance and the effect of efflux pumps on this process.

## METHODS

### Bacterial Strains

We used *E. coli* strains BW25113 (WT), BW25113 Δ*acrR* (AcrAB+), and BW25113 Δ*acrB* (Δ*acrB*) as the parent strains. The WT strain BW25113 is the base strain for the Keio collection (54). For BW25113 *acrR*, we designed primers with homology regions on *acrR* and amplified the kanamycin resistance marker and FRT sites of pKD13 (54). Primers are listed in Table S3. The linear DNA was then treated using a DpnI digest and PCR purification. We electroporated the purified linear DNA into competent BW25113 cells containing the plasmid pSIM6 (64). BW25113 Δ*acrB* was derived from Keio collection strain JW0451 (BW25113 Δ*acrB::kan^R^*) (26). We removed kanamycin resistance markers from BW25113 Δ*acrR::kan^R^* and JW0451 following the pCP20 protocol from Reference (65).

### Determination of MIC

For all experiments, overnight cultures were inoculated from a single colony in 10 mL LB and grown in a 50 mL Erlenmeyer flask at 37°C with 200 rpm orbital shaking. After overnight growth, the optical density at 600 nm (OD_600_) was measured, and the initial volume was diluted back to OD_600_ = 0.1. To determine the MIC of the parent strains (Figure S2), we added a final concentration of 0, 0.2, 0.5, 1, 2, 4, 8, or 12 μg/mL chloramphenicol to each culture; to determine the MIC of the evolved strains (Figure S5), we added 0, 0.5, 1, 2, 5, 10, 20, or 50 μg/mL to each culture. Chloramphenicol stocks were prepared with 100% ethanol. The samples were sealed with evaporation-limiting membranes (Thermo Scientific AB-0580) and grown in 24-well plates at 37°C with 200 rpm orbital shaking. OD_600_ readings were taken using a BioTek Synergy H1m plate reader before incubation (t = 0 h) and after antibiotic exposure (t = 24 h). All experiments were performed in triplicate using biological replicates.

### Experimental Conditions in the eVOLVER

In the eVOLVER, cultures were inoculated from a single colony in LB at 37°C. A stir bar mixed the cultures on a medium setting, or approximately 1000 rpm (39). The LB was supplemented with the detergent Tween20 (Sigma Aldrich Cat. # P1379) at 0.2% (v/v) to reduce spurious OD_600_ measurements caused by biofilm growth on the flask. As Tween20 is a detergent and a potential substrate of the AcrAB-TolC efflux pumps, we also conducted the toxicity curve experiments with Tween20 at our working concentration 0.2% (v/v). We found there was no significant change in resistance for any of the strains in the presence of Tween20 (Figure S6 & Table S4).

Cells were inoculated in the eVOLVER overnight (t » −16 – −14 h) prior to the beginning of the experiment (t = 0 h) to establish steady-state exponential growth. We set the eVOLVER using an upper OD_600_ bound of 0.2 and a lower bound of 0.1; thus, cultures were grown to a turbidity of 0.2 and then diluted back to 0.1 to maintain the turbidostat at approximately constant cell density. Samples were collected during the experiment at set time points (t = 0, 1, 3, 6, 12, 24, 48, and 72 h) and used for downstream analysis. All experiments were performed in triplicate using biological replicates.

At t = 0 h, we introduced chloramphenicol at a predetermined final treatment concentration (0, 0.2, 0.5, 1, 2, 5, 10, or 20 μg/mL). This introduction was implemented by switching the media source from one containing 0 μg/mL chloramphenicol to another containing the final treatment concentration; in addition, we spiked the samples directly with the treatment concentration of chloramphenicol at the same time to avoid a delay due to the time required for media cycling in the turbidostat.

### Downstream Assays and Data Collection from eVOLVER Samples

#### Growth Rate Measurements

Growth rate measurements were calculated after each dilution event using:

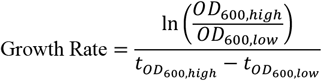

The growth rate between each dilution was then averaged across sampling time points to compare against disc diffusion assays and spot assays. For example, the growth rate given at t = 0 h is the growth rate from t = −6 h to t = 0 h. To evaluate statistically significant differences in growth rate between two time points, we used the paired-*t* test; to evaluate statistically significant differences in growth rate between two strains, we used the *t* test (Table S1A).

#### Antibiotic Disc Diffusion Assay

We aliquoted samples from the eVOLVER, where the OD_600_ from each sample was between 0.1 and 0.2. We used cotton swabs to cover LB agar plates with a layer of the sample (66). An antibiotic disc containing chloramphenicol (30 g) (Thermo Fisher Scientific Cat. # CT0013B) was then placed on the plate. The plate was incubated for 24 h at 37°C. The diameter of the zone of inhibition around each disc was then measured. Diameter of inhibition zones were classified as susceptible, intermediate, or resistant based on Reference (44). Additionally, we calculated the MIC using a mapping between MIC and diameter of inhibition zones for our samples (Figure S5) (67). To evaluate statistically significant differences in diameter of inhibition zones or resistance between two time points, we used the paired-*t* test; to evaluate statistically significant differences in resistance between two genotypes, we used the *t* test (Table S1B).

#### Spot Assay

The samples from the eVOLVER experiment were diluted in PBS in the following dilution series: 1, 10^-1^, 10^-2^, 10^-3^, 10^-4^, and 10^-5^. We then plated 2.5 μL of each dilution on LB agar plates containing 0, 0.5, 1, 2, 5, 10, and 20 μg/mL chloramphenicol. The plates were then incubated for 24 h at 37°C. To count colonies, we identified the dilution factor with the most countable colonies, and recorded the number of colony forming units (*CFU*) and dilution factor (*d*). The CFU/mL for each sample was then calculated by:

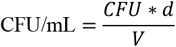

where *V* is the volume plated. We also calculated the proportion of the population able to grow on different concentrations of chloramphenicol by calculating the CFU/mL from LB agar plates containing 0, 0.5, 1, 2, 5, 10, and 20 μg/mL chloramphenicol.

### Whole Genome Sequencing

DNA was extracted from single isolates and parent strains using the QIAGEN DNeasy PowerBiofilm kit. For each strain, we selected three isolates to sequence; each of these isolates originated from a different biological replicate that was evolved under the same experimental conditions (i.e. each isolate comes from a different eVOLVER culture). Samples were sequenced at the Microbial Genome Sequencing Center (MiGS) in Pittsburg, PA, USA, who conducted library preparation and multiplexing using the Illumina Nextera kit series and then sequenced using a NextSeq 550 platform with 150 bp paired-ends and an average coverage of 50 reads. We analyzed reads using *breseq* (68) version 0.35.1. Reads were aligned to the BW25113 Keio reference genome (Accession no.: CP009273) in consensus mode. The treatment concentrations and isolation concentrations used to select each isolate are listed in Table S2.

## Data Availability

Whole genome sequencing data for the parent strains and the isolates are available on GenBank (BioProject: PRJNA666010; Accession no.: CP062239 to CP062250). Other datasets generated during this study are available from the corresponding author upon request.

## ACKNOWLEDGEMENTS

This research was supported by the National Science Foundation (grant No. 1347635) and the National Institutes of Health (grant No. R01AI102922).

We thank Dr. Mo Khalil, Dr. Brandon Wong, Dr. Chris Mancuso, and Zack Heins for their assistance and support with the eVOLVER.

## SUPPLEMENTARY FIGURE CAPTIONS

**Figure S1. Growth rates for each biological replicate and antibiotic concentration.** Mean growth rates for n = 3 biological replicates. Shaded error bars show standard deviation. Cultures grown without chloramphenicol occasionally accumulated biofilms, leading to the large variations in the growth measurements for the 0 μg/ml case.

**Figure S2. Toxicity curves for each parent strain.** Final OD_600_ was measured after 24 h (see Methods: Determination of MIC). Data points show mean values from n = 3 biological replicates, error bars show standard deviation.

**Figure S3. Inhibition zone diameters for each biological replicate and antibiotic concentration.** Mean diameter of inhibition zones (D_inh_) for n = 3 biological replicates. Shaded error bars show standard deviation.

**Figure S4. Colony forming units (CFU) per mL counts for each treatment.** Mean CFU/mL values from n = 3 biological replicates are shown, with error bars denoting standard deviation.

**Figure S5. Linear map between the natural log of the MIC and inhibition zone areas.** Data are from inhibition zone diameters and MIC_90_ for each parent strains (e.g. AcrAB+) and the evolved isolates of each parent strain from three different eVOLVER experiments (e.g. eAcrAB+1, eAcrAB+2, and eAcrAB+3). MIC_90_ is defined as the point where OD_600_ is reduced to 10% of normal growth after 24 h (Figure S2). To find the linear correlation, we calculated the natural log of the MIC_90_ and the inhibition zone areas. The parameters for this map are Q = 30 μg, k = 57.8, and K = −0.971, following the notation from Reference (67).

**Figure S6. Toxicity curves in the presence of Tween20.** Strains grown with and without 0.2% (v/v) Tween20. Data points show mean values from n = 3 biological replicates, error bars show standard deviation.

## SUPPLEMENTARY TABLE CAPTIONS

**Table S1.** P-values from the (paired) *t*-test for quantifying significant differences in **(A)** growth rate or **(B)** resistance as measured by the diameter of inhibition zone between (1) two sequential time points, (2) a given time point and the initial time point, or (3) two strains at a given time point and treatment.

**Table S2.** Summary of sequencing results. Non-clonal mutations for each resistant isolate from eVOLVER experiments. Each isolate from each parent strain is derived from a different biological replicate. In addition to the mutations, the table also lists the treatment concentrations that each isolate evolved at, as well as the concentration of chloramphenicol that the isolate was selected on at t = 72 h. Genetic regions that do not exist in the parent strain are grayed out.

**Table S3.** Primers containing 40-nt homology regions for *acrR* knockout. Bold letters denote the active priming region to amplify pKD13 from Reference (54).

**Table S4.** P-values from the paired *t*-test to assess statistically significant differences in growth between samples treated with Tween20 at 0.0% and 0.2% (v/v) as shown in Figure S6.

