## Supplementary figures and images for "Mapping the Role of AcrAB-TolC Efflux Pumps in the Evolution of Antibiotic Resistance Reveals Near-MIC Treatments Facilitate Resistance Acquisition"

### Figure S1

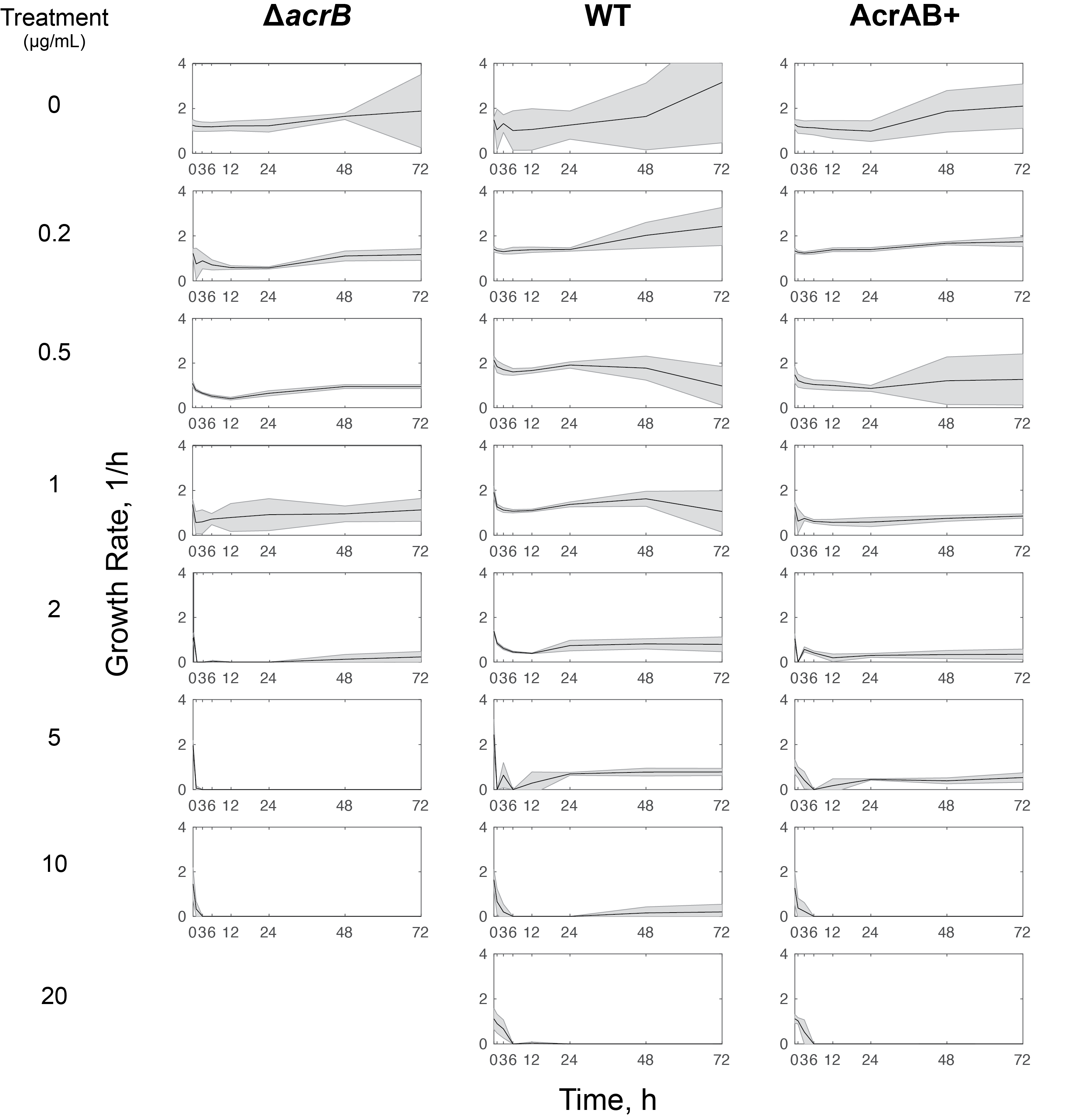

### Figure S2

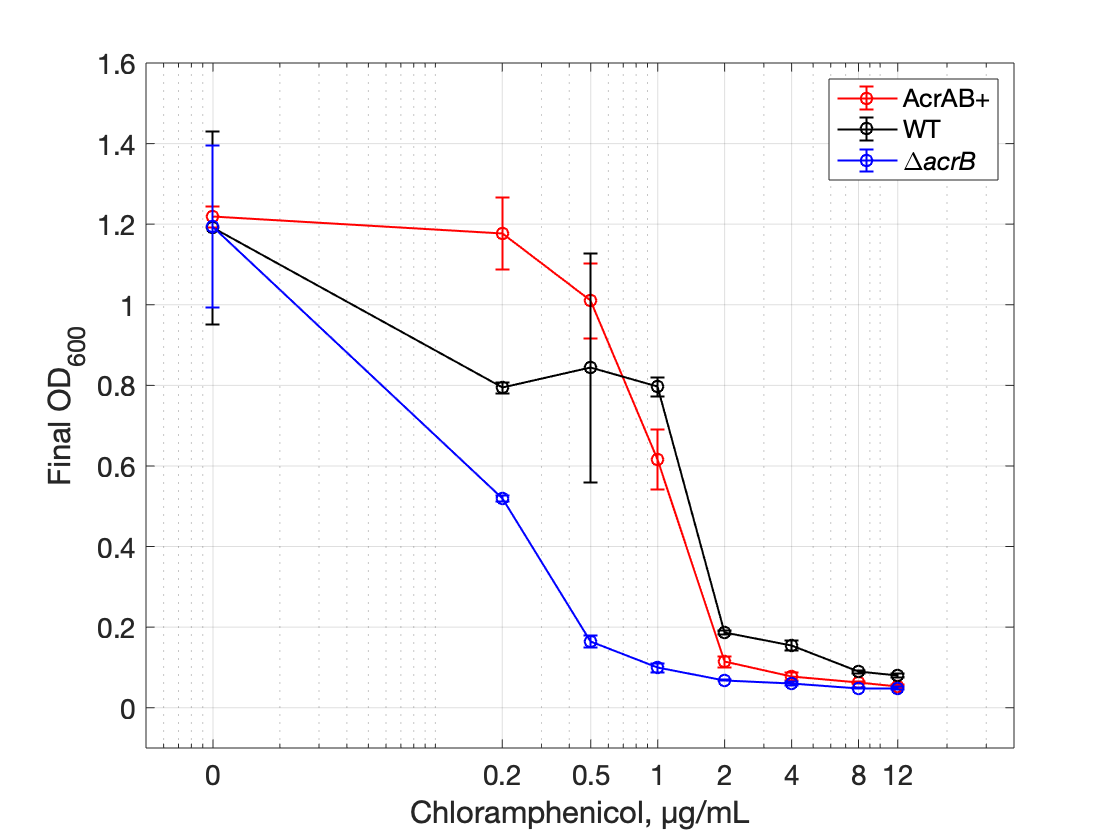

### Figure S3

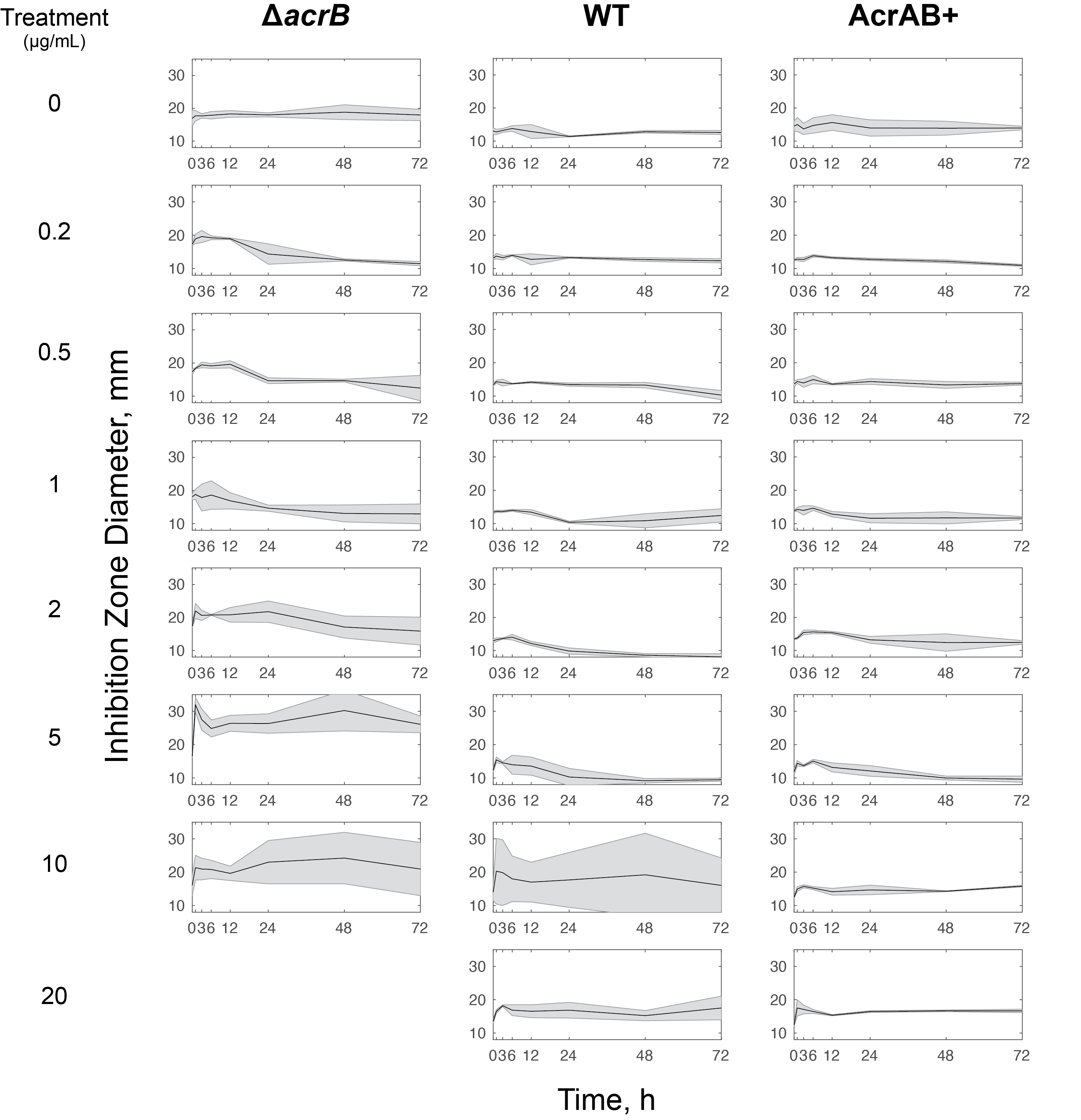

### Figure S4

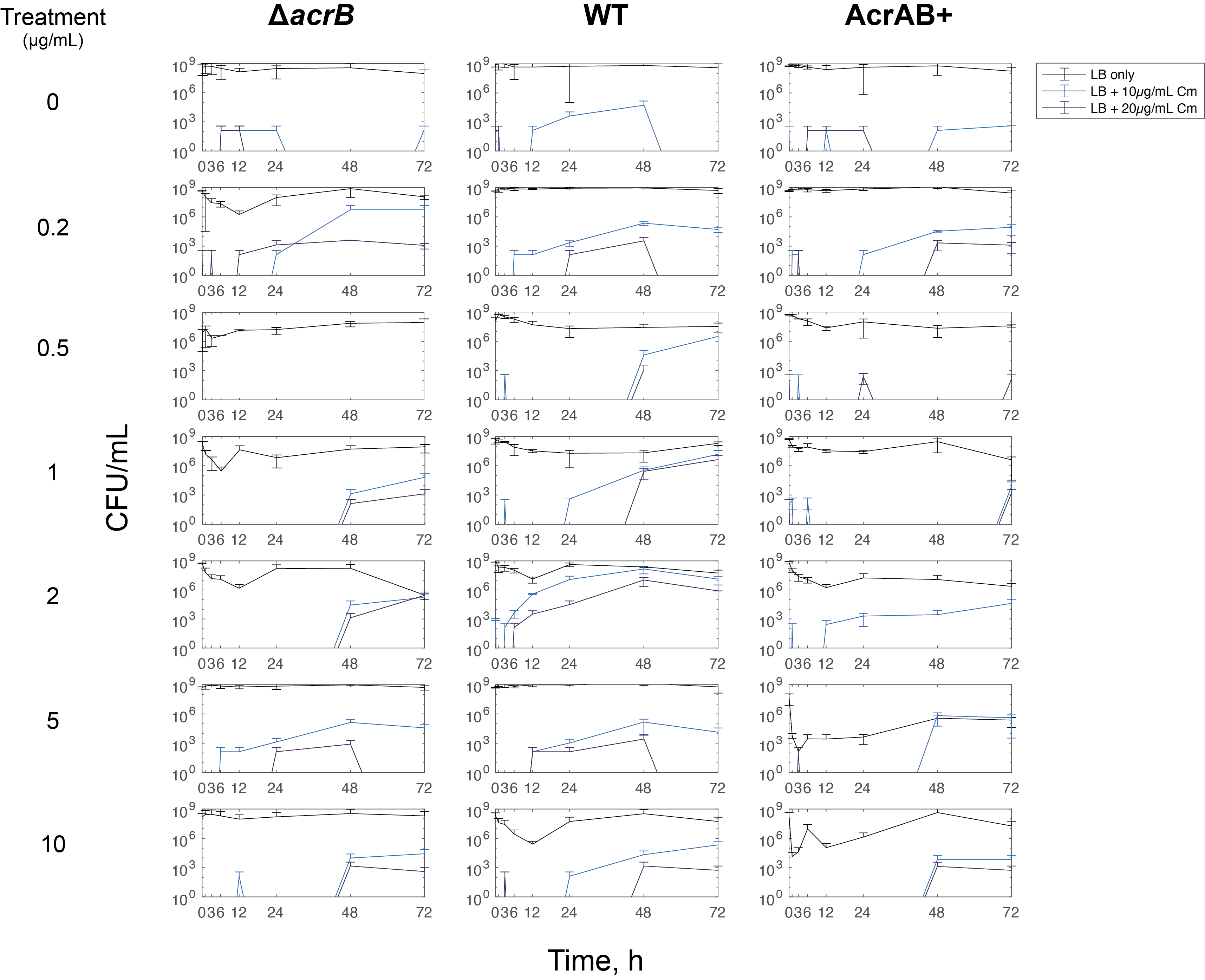

### Figure S5

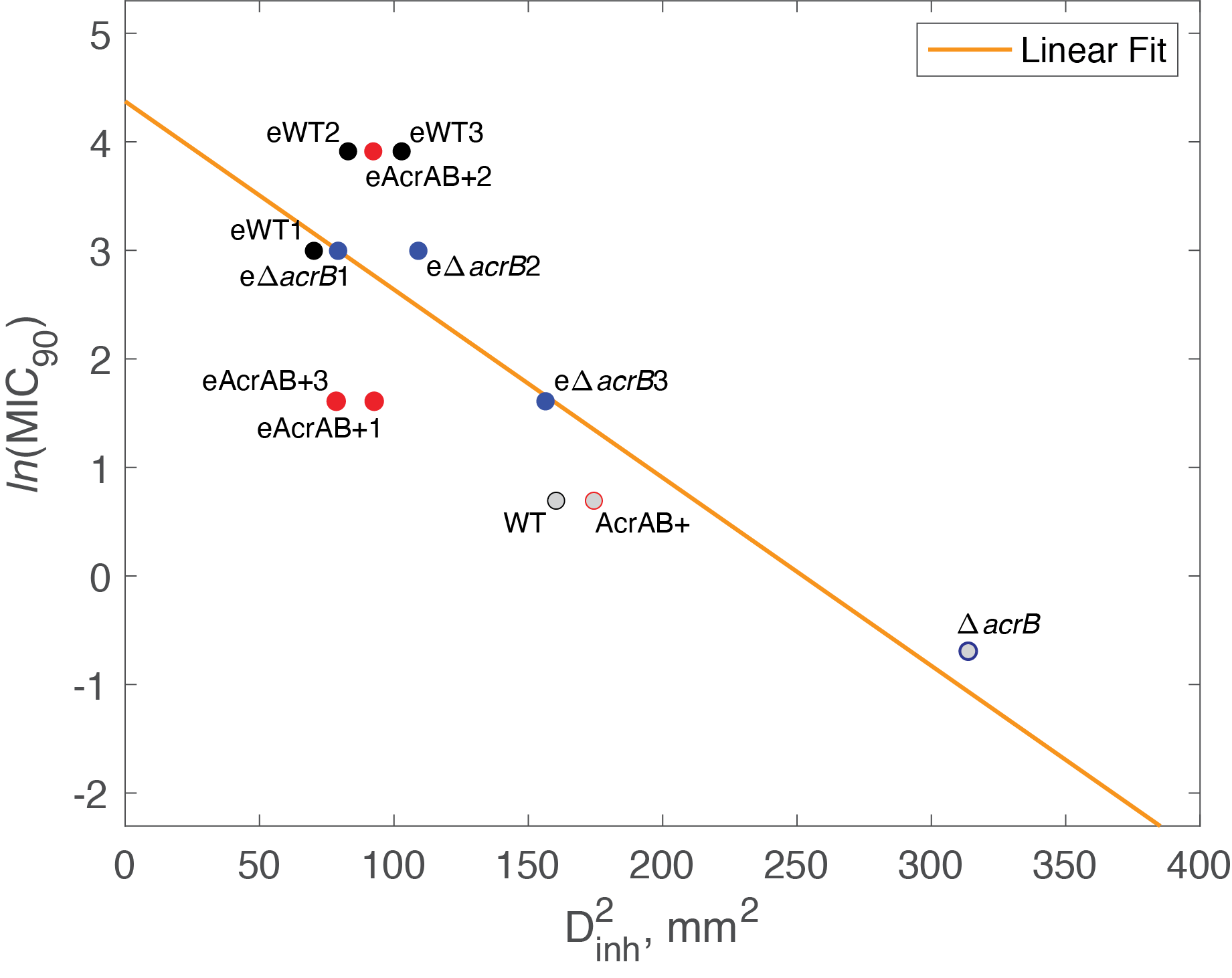

### Figure S6

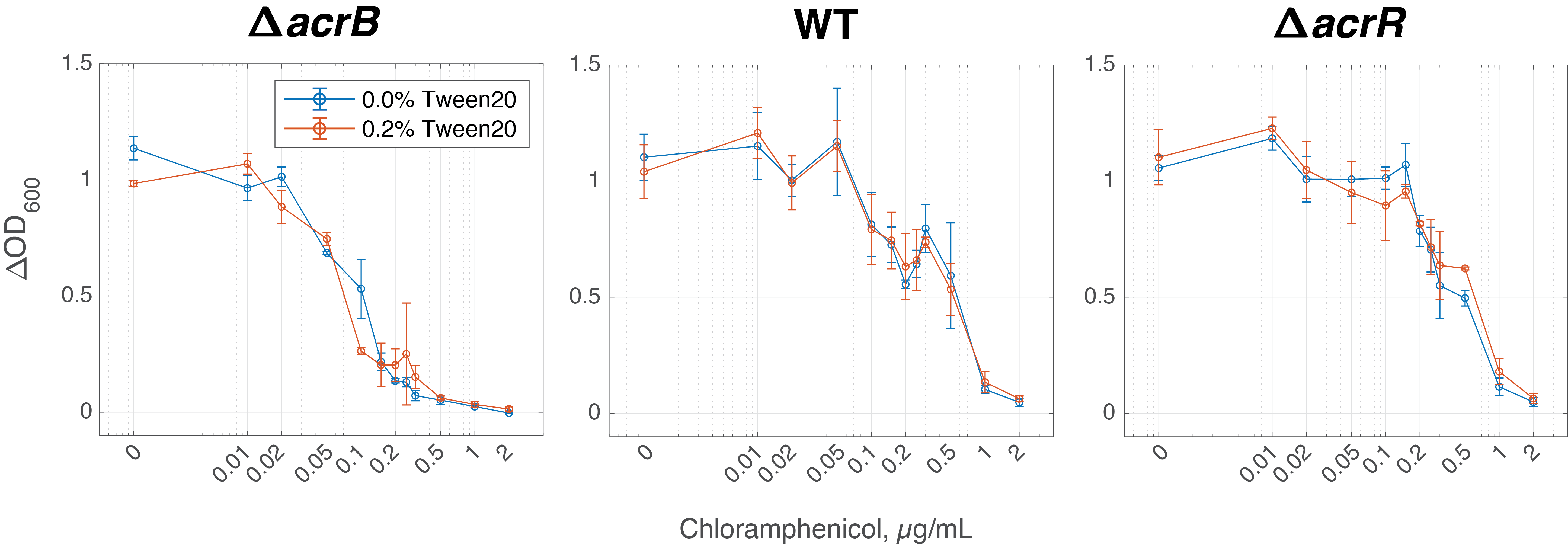
