## Supplementary material for "Mapping the Role of AcrAB-TolC Efflux Pumps in the Evolution of Antibiotic Resistance Reveals Near-MIC Treatments Facilitate Resistance Acquisition": Table S2

| Parent Strain | | | **WT** | | | **AcrAB+** | | | **Δ*acrB*** | | |
| --- | --- | --- | --- | --- | --- | --- | --- | --- | --- | --- | --- |
| Treatment Concentration | | | **2 µg/mL Cm** | | | **5 µg/mL Cm** | | | **1 µg/mL Cm** | | |
| **Region** | **Mutation** | **Position** | 1 | 2 | 3 | 1 | 2 | 3 | 1 | 2 | 3 |
| ***acrR*** | IS1 + 4bp | 481,420 |  |  | X |  |  |  |  |  |  |
|  | IS5 + 8bp | 481,481 | X |  |  |  |  |  |  |  |  |
| **P*_acrRAB_*** | IS2 + 4bp | 481,163 |  |  |  |  |  | X |  |  |  |
|  | Δ 1bp | 481,174 |  |  |  | X |  |  |  |  |  |
|  | T🡪C | 481,187 |  |  |  |  | X |  |  |  |  |
| ***acrB*** | Q569L | 478,154 |  | X |  |  |  | X |  |  |  |
|  | V139F | 479,445 |  |  |  |  | X |  |  |  |  |
| ***marR*** | + 1bp | 1,613,590 |  | X |  |  |  |  |  |  |  |
|  | T72P | 1,613,590 | X |  |  |  |  |  |  |  |  |
|  | V84E | 1,613,267 |  |  | X |  |  |  |  |  |  |
| ***acrS*** | IS5 + 4bp | 3,407,126 |  |  |  |  |  |  | X | X |  |
|  | IS2 + 4bp | 3,407,133 |  |  |  |  |  |  |  |  | X |
| ***rpoB*** | K126Q | 4,174,956 |  |  |  | X |  |  |  |  |  |
| ***fimD*** | T393N | 4,536,090 |  |  |  |  | X |  |  |  |  |
| ***yhjB*** | IS4 + 12bp | 3,664,650 |  |  |  |  |  | X |  |  |  |
| ***clpX*** | IS186 + 2bp | 454,251 |  |  |  |  |  | X |  |  |  |
| ***selA*** | D441G | 3,753,288 |  |  |  |  |  |  | X |  |  |
| ***rrsG*** | +58 bp | 2,723,638 |  |  |  |  |  |  | X |  |  |
| Isolation [Cm] (µg/mL) | | | 20 | 20 | 20 | 10 | 10 | 10 | 10 | 5 | 5 |
