## Supplementary material for "Mapping the Role of AcrAB-TolC Efflux Pumps in the Evolution of Antibiotic Resistance Reveals Near-MIC Treatments Facilitate Resistance Acquisition": Table S3

| **Direction** | **Primer (5’ to 3’)** |
| --- | --- |
| Forward | ATGTATGTAAATCTAACGCCTGTAAATTCACGAACATATG**GTGTAGGCTGGAGCTGCTTC** |
| Reverse | CCTGGAGTCAGATTCAGGGTTATTCGTTAGTGGCAGGATT**GATCCGTCGACCTGCAGTT** |
