## Supplementary material for "Mapping the Role of AcrAB-TolC Efflux Pumps in the Evolution of Antibiotic Resistance Reveals Near-MIC Treatments Facilitate Resistance Acquisition": Table S4

|  | **Chloramphenicol (µg/mL)** | | | | | | | | | | | |
| --- | --- | --- | --- | --- | --- | --- | --- | --- | --- | --- | --- | --- |
|  | **0** | **0.01** | **0.02** | **0.05** | **0.10** | **0.15** | **0.20** | **0.25** | **0.30** | **0.50** | **1** | **2** |
| **WT** | 0.389 | 0.355 | 0.919 | 0.919 | 0.900 | 0.567 | 0.494 | 0.832 | 0.474 | 0.714 | 0.275 | 0.107 |
| **AcrAB+** | 0.375 | 0.510 | 0.586 | 0.367 | 0.225 | 0.236 | 0.446 | 0.938 | 0.435 | 0.016 | 0.039 | 0.053 |
| **Δ*acrB*** | 0.032 | 0.138 | 0.142 | 0.089 | 0.053 | 0.809 | 0.229 | 0.429 | 0.187 | 0.273 | 0.430 | 0.058 |
